# Transcriptome from saffron (*Crocus sativus*) plants in Jammu and Kashmir reveals abundant soybean mosaic virus transcripts and several putative pathogen bacterial and fungal genera

**DOI:** 10.1101/079186

**Authors:** Sandeep Chakraborty

## Abstract

Saffron (*Crocus sativus*) is a spice with immense economic and medicinal relevance, due to its anticancer and chemopreventive properties. Although the genomic sequence of saffron is not publicly available, the RNA-seq based transcriptome of saffron from Jammu and Kashmir provides several, yet explored, insights into the metagenome of the plant from that region. In the current work, sequence databases were created in the YeATS suite from the NCBI and Ensembl databases to enable faster comparisons. These were used to determine the metagenome of saffron. Soybean mosaic virus, a potyvirus, was found to be abundantly expressed in all five tissues analyzed. Recent studies have highlighted that issues arising from latent potyvirus infections in saffron is severely underestimated. Bacterial and fungal identification is made complex due to symbiogenesis, especially in the absence of the endogenous genome. Symbiogenesis results in transcripts having significant homology to bacterial genomes and eukaryotic genomes. A stringent criterion based on homology comparison was used to identify bacterial and fungal transcripts, and inferences were constrained to the genus level. *Leifsonia*, *Elizabethkingia* and *Staphylococcus* were some of the identified bacteria, while *Mycosphaerella* and *Pyrenophora* were among the fungi detected. Among the bacterial genera, *L. xyli* is the causal agent for ratoon stunting disease in sugarcane, while *E. meningoseptica* and *S. haemolyticus*, having acquired multiresistance against available antimicrobial agents, are important in clinical settings. *Mycosphaerella* and *Pyrenophora* incorporate several pathogenic species. It is shown that a transcript from heat shock protein of the fungi *Cladosporium cladosporioides* has been erroneously annotated as a saffron gene. The detection of these pathogens should enable proper strategies for ensuring better yields. The functional annotation of proteins in the absence of a genome is subject to errors due to the existence of significantly homologous proteins in organisms from different branches of life.

## Introduction

Saffron, possibly one of the most expensive spices, is an anticancer and chemopreventive agent [1, 2], and is also effective for treating major depressive disorder symptoms [3, 4]. The genome of saffron is not publicly available, although http://www.crocusgenome.org/ provides a online interface to query individual sequences. The RNA-seq transcriptome [5, 6] of saffron was independently sequenced by two research groups in order to gain deeper insights into the genes involved in apocarotenoid biosynthesis [7, 8]. Previously, the YeATS suite identified several artifacts arising from RNA-seq assembly [9, 10], and has been used to analyze the walnut transcriptome revealing the biodiversity and plantmicrobe interactions in twenty different tissues from walnut in California [11].

A BLAST to the complete ’nt/nr’ database, as done previously, is not time efficient [9–11]. In the current work, the NCBI and Ensembl database was used to create smaller, yet comprehensive, databases for viruses (V-DB), bacterial (B-DB), fungal (F-DB) and plants (mitochondria, chloroplast and ribosomes – CMR-DB), and were added to the YeATS suite for a time efficient query of the metagenome. In the absence of a genome, identification of saffron derived transcripts is not straightforward. Using these databases, the metagenome of saffron from Jammu and Kashmir was derived from the transcriptome of saffron. http://nipgr.res.in/mjain.html?page=saffron provided the ABySS [12] assembled transcriptome with 105269 contigs. Mosaic virus derived transcripts were found to be abundantly expressed in all five tissues analyzed [13]. Mosaic viruses can be responsible for significant yield losses [14]. Key organelles of eukaryotes, like the mitochondrion and chloroplast, possibly originated as a symbiosis between distinct single-celled organisms [15]. Symbiogenesis results in transcripts that have significant homology to both bacterial and eukaryotic genomes. Consequently, a stringent constraint that compared homology scores was used to choose bacterial and fungal transcripts. *Leifsonia*, *Elizabethkingia* and *Staphylococcus* were some of the identified bacteria, while *Mycosphaerella* and *Pyrenophora* were among the fungi detected. These bacterial and fungal genera incorporate several pathogenic species. A proper disease-management strategy can be devised based on this knowledge. It is also shown that protein-based annotation, based on open reading frames of transcripts, can be error prone due to the existence of significantly homologous proteins in organisms from different branches of life, like fungi, plants and viruses.

## Materials and methods

### Obtaining transcriptome and expression matrix from different tissues of saffron

http://nipgr.res.in/mjain.html?page=saffron provided the ABySS [12] assembled transcriptome with 105269 contigs. The site provides the sequences (Saffron transcriptome assembly.fa), annotation (Saffron transcriptome assembly.fa) and expression values for different tissues (Saffron expression matrix).

### Obtaining the viral sequences

“http://www.ncbi.nlm.nih.gov/genomes/GenomesGroup.cgi?taxid=10239” provides a retrieve option to get all virus genomes (viral.nt.fa:n=7331 in Dataset1). For viral protein sequences, the NCBI database was queried for “Viruses, RefSeq” (viral.nr.refseq.fa, n=50021 in Dataset1)

### Obtaining the bacterial genomes

“ftp://ftp.ncbi.nlm.nih.gov/genomes/refseq/bacteria/assemblysummary.txt” provides details of available bacteria genomes. This was parsed, and the first occuring species of a genus was chosen randomly (getBacterialGenomes.csh:n=1355 in Dataset1).

### Obtaining plant mitochondrial, chloroplast and ribosomal sequences

The NCBI database was queried for : Plants, RefSeq, Mitochondrion/Chloroplast/Ribosomal rRNA/Ribosomal mRNA. These were combined in a single file (DB.list CHLORO MITO MRIBO RRIBO, n=40049 in Dataset1).

### Obtaining plant mitochondrial, chloroplast and ribosomal sequences

The fungal sequences were obtained from the Ensembl site [16]. A random species was chosen for each genus (list.ensembl.fungi.txt in Dataset1, n=222).

The YeATS suite was used extensively to query these databases using the BLAST command-line interface [17]. The BLAST bitscore was used as a comparison metric instead of the Evalue since it allows differentiation for high homologies where Evalue goes to zero.

## Results and discussion

### Viral transcripts

Transcripts that map to virus genomes (V-DB) clearly demonstrates the presence of the several mosaic virus (MV) species (Table 1). MV are not a single taxon, although they are all ssRNA positive-strand viruses [13]. For example, tobacco MV is a Tobamovirus, while all other MVs in Table 1 are Potyviruses. Previously, saffron has been shown to be infected with bean yellow MV [18] and Narcissus MV [19]. *Iris*, which belong to the same *Iridaceae* family as saffron, was shown to be infected with *Iris* severe MV [20,21] and Narcissus MV [22]. A recent study highlights that the threat from these MVs is seriously underestimated [23].

**Table 1:** Transcripts mapping to viruses sorted based on BLAST bitscore (BBS): There are several transcripts which match to mosaic viruses (MV). Only CSTC028935 has a significant match (gb EX144186.1) to the unpublished saffron genome, queried through the online interface (http://www.crocusgenome.org/?pageid=20). It is very unlikely that the transcript is a part of the saffron genome since it encodes an open reading frame (1587 amino acid long) with a very significant match (79% identity) to the soybean MV polyprotein (Accid:ADK60773.1).

| TranscriptID | Accession ID | Description | BBS |
| --- | --- | --- | --- |
| CSTC028935 | NC_003397.1. | Bean common MV | 1328 |
| CSTC098263 | NC_001367.1. | Tobacco MV | 952 |
| CSTC029938 | NC_001367.1. | Tobacco MV | 876 |
| CSTC042852 | NC_007216.1. | Wisteria vein MV | 651 |
| CSTC006658 | NC_007216.1. | Wisteria vein MV | 608 |
| CSTC103119 | NC_001367.1. | Tobacco MV | 553 |
| CSTC003939 | NC_007216.1. | Wisteria vein MV | 459 |
| CSTC045789 | NC_009742.1. | Telosma MV | 176 |

The complete list of virus genome matches with a BLAST bitscore (BBS) cutoff=150 (~Evalue=1E-30) (see viral.anno.nucleotide.txt in Dataset1, n=8). Transcripts with the highest significance (Table 1) were BLAST’ed through the online interface to the saffron genome. Only CSTC028935 had a significant match to ”gb EX144186.1 EX144186 cr28 F19 Saffron (*Crocus sativus*) mature stigma lambda Unizap”, with a 97% identity, although it is unlikely that this is part of the saffron genome. CSTC028935 encodes a 1587 long open reading frame (ORF) that is 79% identical to the soybean MV polyprotein (Accid:ADK60773.1). Expectedly, none of these viral transcripts were annotated previously in Saffron functional annotation [7].

### Bacterial transcripts

The endosymbiont theory posits that key organelles of eukaryotes, like the mitochondrion and chloroplast, originated as a symbiosis between distinct single-celled organisms [15]. While the origin of this symbiosis remains a subject of debate, the fact that prokaryotes and eukaryotes share significant homology in their genomes is established beyond doubt [24]. This presents a certain degree of uncertainty in identifying the metagenome through nucleotide sequencing, especially when the genome of the endogenous organism is not known. This uncertainty is demonstrated in the analysis below.

First, the saffron transcripts were BLAST’ed to the locally created bacterial genome database (B-DB) (1355 bacterial genomes, see Methods). There were 430 matching transcripts (BBS=200, Evalue cutoff=~1E-50, see BACT.200.anno.sort in Dataset1). A combined database comprising chloroplast, mitochondrial and ribosomal genes (CMR-DB) using a more relaxed Evalue constraint gave 573 matching transcripts (BBS=60, Evalue=1E-10, CHLOROMITORIBO.60.anno.sort in Dataset1). Since the organism under focus here is a plant, the Evalue is more relaxed for these matches. In case a transcript matched both B-DB and CMR-DB, the percentage difference in the BBS was used as a metric to select bacterial genes. There about 150 transcripts where the B-DB BBS score was 60% greater than than the CMR-DB score. About 80 transcripts had no match in CMR-DB. Combined with the high cutoff required for bacterial identification, and low threshold for CMR-DB assignment, these 230 transcripts can be assigned to bacterial species (BACT.MITOCHLORO.anno.sort in Dataset1).

Bacterial transcripts sorted based on BBS revealed the presence of several bacterial genera (Table 2). *Enterococcus* is a large genus of lactic acid bacteria, of which *E. faecalis* and *E. faecium* are common commensal organisms in the intestines of humans. *E. silesiacus* was found in two water isolates [25]. Similarly, *Staphylococcus haemolyticus* is the most frequent aetiological factors of staphylococcal infections, attributed to its ability to acquire multiresistance against available antimicrobial agents [26]. *Leifsonia xyli* is the causal agent of ratoon stunting disease in sugarcane [27], and is surprisingly genetically conserved in isolates from different countries [28]. The commmonly occuring gram-negative bacillus *Elizabethkingiaa* is found in soil, river water and reservoirs. *Elizabethkingia meningoseptica* is resistant to most antibiotics used in the intensive care setting [29], and has been responsible for an outbreak arising from tap contamination [30]. A recent case of bacteremia has been attributed to *Elizabethkingia meningoseptica* [31].

**Table 2:**
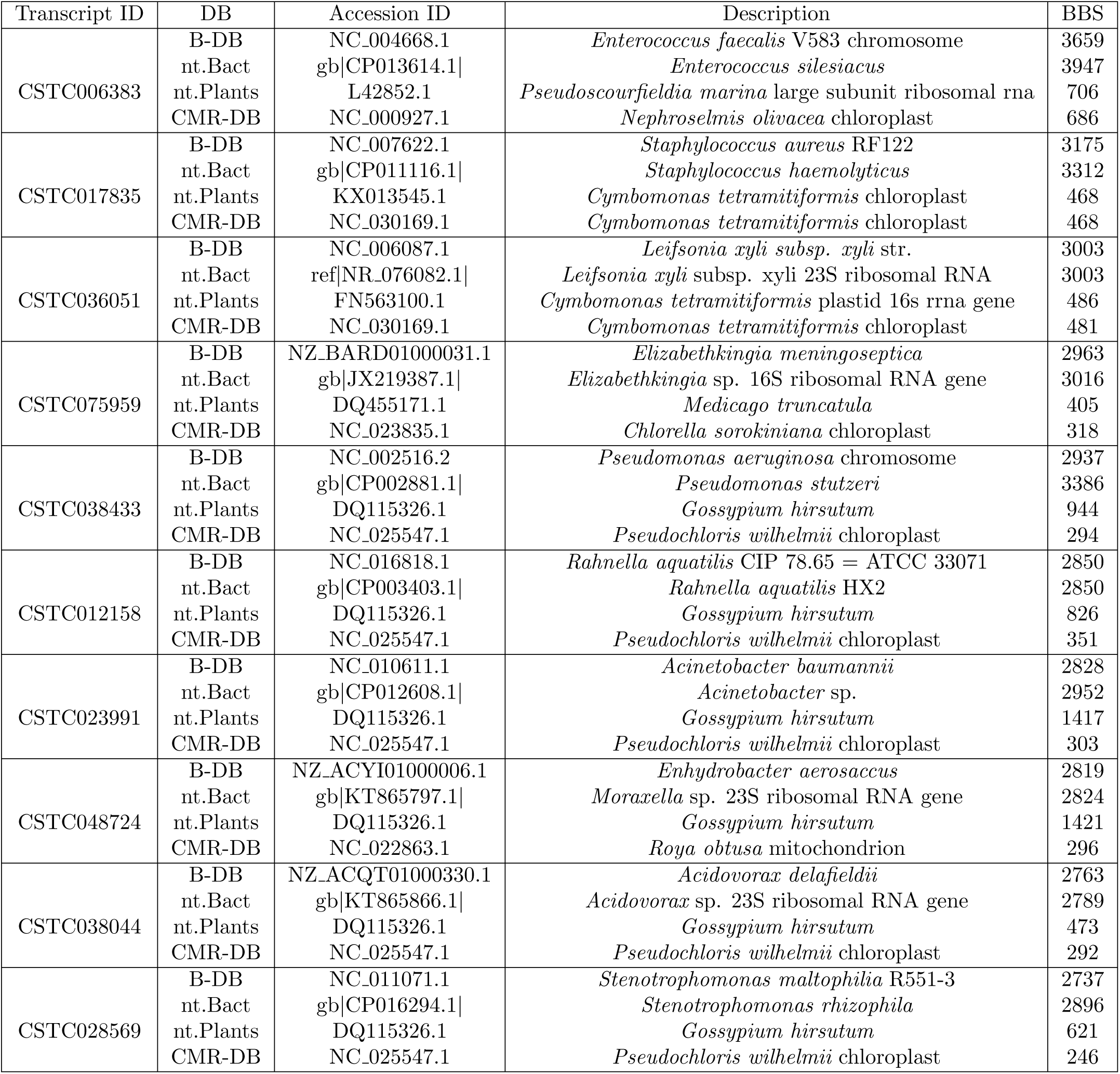
Bacterial transcripts sorted based on BLAST bitscore (BBS): These were identified by comparing the safffron transcriptome to the local bacterial database (B-DB). The homology of these bacterial transcripts to the CMR-DB (Chloroplast-Mitochondrion-Ribosome-DB) provides support for the endosymbiont theory. The transcripts matching the bacterial database were subsequently BLAST’ed to the complete ’nt’ database and the ’plant nt’ database. Although, these bacterial transcripts also have significant matches to plant genomes, they are significantly lower than the matches to bacterial genomes. Interestingly, apart from the only mitochondrional transcript CSTC048724 which had a different genus (*Enhydrobacter* vs *Moraxella*), all other where chloroplast transcripts, and had the same genus when compared to the selected bacterial database versus the complete ’nt’ database. As expected, none of these transcripts were annotated previously [7].

| Transcript ID | DB | Accession ID | Description | BBS |
| --- | --- | --- | --- | --- |
| CSTC006383 | B-DB | NC_004668.1 | *Enterococcus faecalis* V583 chromosome | 3659 |
|  | nt.Bact | gb\|CP013614.1\| | *Enterococcus silesiacus* | 3947 |
|  | nt.Plants | L42852.1 | *Pseudoscourfieldia marina* large subunit ribosomal rna | 706 |
|  | CMR-DB | NC_000927.1 | *Nephroselmis olivacea* chloroplast | 686 |
| CSTC017835 | B-DB | NC_007622.1 | *Staphylococcus aureus* RF122 | 3175 |
|  | nt.Bact | gb\|CP011116.1\| | *Staphylococcus haemolyticus* | 3312 |
|  | nt.Plants | KX013545.1 | *Cymbomonas tetramitiformis* chloroplast | 468 |
|  | CMR-DB | NC_030169.1 | *Cymbomonas tetramitiformis* chloroplast | 468 |
| CSTC036051 | B-DB | NC_006087.1 | *Leifsonia xyli* subsp. *xyli* str. | 3003 |
|  | nt.Bact | ref\|NR_076082.1\| | *Leifsonia xyli* subsp. *xyli* 23S ribosomal RNA | 3003 |
|  | nt.Plants | FN563100.1 | *Cymbomonas tetramitiformis* plastid 16s rRNA gene | 486 |
|  | CMR-DB | NC_030169.1 | *Cymbomonas tetramitiformis* chloroplast | 481 |
| CSTC075959 | B-DB | NZ_BARD01000031.1 | *Elizabethkingia meningoseptica* | 2963 |
|  | nt.Bact | gb\|JX219387.1\| | *Elizabethkingia* sp. 16S ribosomal RNA gene | 3016 |
|  | nt.Plants | DQ455171.1 | *Medicago truncatula* | 405 |
|  | CMR-DB | NC_023835.1 | *Chlorella sorokiniana* chloroplast | 318 |
| CSTC038433 | B-DB | NC_002516.2 | *Pseudomonas aeruginosa* chromosome | 2937 |
|  | nt.Bact | gb\|CP002881.1\| | *Pseudomonas stutzeri* | 3386 |
|  | nt.Plants | DQ115326.1 | *Gossypium hirsutum* | 944 |
|  | CMR-DB | NC_025547.1 | *Pseudochloris wilhelmii* chloroplast | 294 |
| CSTC012158 | B-DB | NC_016818.1 | *Rahnella aquatilis* CIP 78.65 = ATCC 33071 | 2850 |
|  | nt.Bact | gb\|CP003403.1\| | *Rahnella aquatilis* HX2 | 2850 |
|  | nt.Plants | DQ115326.1 | *Gossypium hirsutum* | 826 |
|  | CMR-DB | NC_025547.1 | *Pseudochloris wilhelmii* chloroplast | 351 |
| CSTC023991 | B-DB | NC_010611.1 | *Acinetobacter baumannii* | 2828 |
|  | nt.Bact | gb\|CP012608.1\| | *Acinetobacter* sp. | 2952 |
|  | nt.Plants | DQ115326.1 | *Gossypium hirsutum* | 1417 |
|  | CMR-DB | NC_025547.1 | *Pseudochloris wilhelmii* chloroplast | 303 |
| CSTC048724 | B-DB | NZ_ACYI01000006.1 | *Enhydrobacter aerosaccus* | 2819 |
|  | nt.Bact | gb\|KT865797.1\| | *Moraxella* sp. 23S ribosomal RNA gene | 2824 |
|  | nt.Plants | DQ115326.1 | *Gossypium hirsutum* | 1421 |
|  | CMR-DB | NC_022863.1 | *Roya obtusa* mitochondrion | 296 |
| CSTC038044 | B-DB | NZ_ACQT01000330.1 | *Acidovorax delafieldii* | 2763 |
|  | nt.Bact | gb\|KT865866.1\| | *Acidovorax* sp. 23S ribosomal RNA gene | 2789 |
|  | nt.Plants | DQ115326.1 | *Gossypium hirsutum* | 473 |
|  | CMR-DB | NC_025547.1 | *Pseudochloris wilhelmii* chloroplast | 292 |
| CSTC028569 | B-DB | NC_011071.1 | *Stenotrophomonas maltophilia* R551-3 | 2737 |
|  | nt.Bact | gb\|CP016294.1\| | *Stenotrophomonas rhizophila* | 2896 |
|  | nt.Plants | DQ115326.1 | *Gossypium hirsutum* | 621 |
|  | CMR-DB | NC_025547.1 | *Pseudochloris wilhelmii* chloroplast | 246 |

**Table 3:** Counts of transcripts mapping to viruses (V) and bacteria (B) sorted based on BLAST bitscore: Viral transcripts from mosaic viruses are abundantly present in all five tissues analyzed. In comparison, bacterial transcripts are fewer in numbers.

|  | TranscriptID | TrLen | CORM | TEPAL | LEAF | STIGMA | STAMEN |
| --- | --- | --- | --- | --- | --- | --- | --- |
| V | CSTC028935 | 4762 | 167821.9 | 242246.9 | 163341.6 | 1067049.2 | 29630.3 |
|  | CSTC098263 | 781 | 1 | 49.9 | 0 | 0 | 0 |
|  | CSTC029938 | 889 | 0 | 85.4 | 0 | 0 | 0 |
|  | CSTC042852 | 7175 | 78307.1 | 87094.2 | 62997.3 | 289732.9 | 8391.9 |
|  | CSTC006658 | 3519 | 40341.8 | 86292.2 | 48059.2 | 237711.2 | 4193.8 |
|  | CSTC103119 | 759 | 0 | 60.4 | 0 | 0 | 0 |
|  | CSTC003939 | 2758 | 11938.7 | 34691.7 | 8590.5 | 51597 | 1335.8 |
|  | CSTC045789 | 558 | 25667.9 | 8440 | 10437.1 | 69069.3 | 2162.8 |
| B | CSTC006383 | 2924 | 430.7 | 785.7 | 25.7 | 280.2 | 263.7 |
|  | CSTC017835 | 1898 | 247.1 | 221.6 | 38.0 | 139.4 | 8.6 |
|  | CSTC036051 | 2315 | 177.3 | 307.0 | 19.5 | 14.3 | 12.9 |
|  | CSTC075959 | 1885 | 263.8 | 148.7 | 129.3 | 175.8 | 14.3 |
|  | CSTC038433 | 1864 | 896.8 | 196.7 | 70.8 | 220.2 | 30.1 |
|  | CSTC012158 | 1646 | 472.4 | 490.2 | 16.4 | 362.4 | 322.5 |
|  | CSTC023991 | 1780 | 810.2 | 497.9 | 119.1 | 203.7 | 10.0 |
|  | CSTC048724 | 1648 | 199.2 | 251.3 | 32.9 | 52.2 | 21.5 |
|  | CSTC038044 | 1761 | 248.2 | 270.5 | 40.0 | 215.9 | 18.6 |
|  | CSTC028569 | 2054 | 115.8 | 159.2 | 17.4 | 117.2 | 18.6 |

**Table 4:** Transcripts with the highest cumulative counts : Apart from one viral transcript (CSTC028935), and one cytochrome P450 encoding ORF (CSTC042320), all other transcripts are ribosomal RNA or from the chloroplast/mitochondria. These eight transcripts also have significant matches to the bacterial database, although their homology to the plant genomes is significantly higher. The cytochrome P450 transcript is annotated by analyzing the open reading frame, since the nucleotide sequence has no match in the complete ’nt’ database.

| TranscriptID | CORM | TEPAL | LEAF | STIGMA | STAMEN | Description |
| --- | --- | --- | --- | --- | --- | --- |
| CSTC043388 | 1311047.7 | 4516814.8 | 423456.4 | 134222.0 | 1403873.5 | 26S rRNA |
| CSTC048227 | 642066.9 | 2044589.8 | 255631.2 | 77501.7 | 509226.4 | 18S rRNA |
| CSTC031703 | 572899.7 | 1513910.7 | 150241.9 | 40571.2 | 998203.4 | 26S rRNA |
| CSTC009284 | 360533.7 | 1334369.3 | 83657.0 | 23579.3 | 628958.8 | 26S rRNA |
| CSTC028935 | 167821.9 | 242246.9 | 163341.6 | 1067049.2 | 29630.3 | Bean Mosaic Virus |
| CSTC018503 | 212853.0 | 482137.8 | 39453.3 | 13557.8 | 277618.9 | 26S rRNA |
| CSTC042320 | 57.4 | 143.9 | 86.2 | 805965.3 | 9905.4 | Cytochrome P450 (Protein) |
| CSTC044767 | 33477.2 | 42562.8 | 696846.6 | 20970.8 | 14874.6 | Chloroplast |
| CSTC050677 | 50947.8 | 477696.2 | 23367.5 | 12243.9 | 99507.3 | Mitochondria |
| CSTC019287 | 17553.0 | 29246.6 | 467484.7 | 14051.0 | 7976.2 | Chloroplast |

**Table 5:** Transcripts with highest counts – comparing the significance of homology to plant and bacterial genomes: These transcripts are from Table 4, barring a virus derived transcript and the cytochrome P450 gene. They have been BLAST’ed t the complete ’nt’ database. The endosymbiont theory is amply supported by the significant matches of these plant mitochondrial and chloroplast transcripts to bacterial genomes.

|  |  |  |  |
| --- | --- | --- | --- |
| CSTC043388 | AY095471.1 | *Bulbine succulenta* 26S ribosomal RNA gene, partial sequence | 2492 |
| CSTC043388 | NZ_LDOD01000051.1 | *Tenacibaculum mesophilum* strain HMG1 scaffold51, whole genome | 990 |
| CSTC048227 | HM640735.1 | *Hesperocallis undulata* voucher Cranfill Schmid s.n.(JEPS) 18S | 1664 |
| CSTC048227 | NZ_LDOD01000055.1 | *Tenacibaculum mesophilum* strain HMG1 scaffold55, whole genome | 178 |
| CSTC031703 | KM117261.1 | *Wisteria floribunda* 18S ribosomal RNA gene, internal transcribed | 1068 |
| CSTC031703 | NZ_ATXL01000109.1 | *Bavariicoccus seileri* DSM 19936 H541DRAFT_scaffold00109.109_C, | 549 |
| CSTC009284 | AF389262.1 | *Dicentra eximia* 26S ribosomal RNA gene, partial sequence | 1033 |
| CSTC009284 | NZ_ATXL01000109.1 | *Bavariicoccus seileri* DSM 19936 H541DRAFT_scaffold00109.109_C, | 651 |
| CSTC018503 | HG970335.2 | *Fusarium graminearum* chromosome 4, complete genome | 1993 |
| CSTC018503 | NZ_ATXL01000109.1 | *Bavariicoccus seileri* DSM 19936 H541DRAFT_scaffold00109.109_C, | 1509 |
| CSTC044767 | KP899622.1 | *Dioscorea zingiberensis* voucher DZING20150203V3 chloroplast, | 5288 |
| CSTC044767 | NC_006576.1 | *Synechococcus elongatus* PCC 6301 DNA, complete genome | 1999 |
| CSTC050677 | JN375330.1 | *Phoenix dactylifera* mitochondrion, complete genome | 3192 |
| CSTC050677 | NZ_BBVC01000023.1 | *Caedibacter varicaedens* DNA, contig: Cva_contig000023, whole | 638 |
| CSTC019287 | KM014691.1 | *Iris gatesii* voucher T06-52 plastid, complete genome | 6335 |
| CSTC019287 | NZ_GL890965.1 | *Moorea producens* 3L scf52112, whole genome shotgun sequence | 544 |

An example of a transcript with significant homology to bacterial and plant genomes is CSTC008113. This has the best match in the ’nt’ database to *Vicia americana*, a legume known commonly as American vetch and purple vetch, with BBS=957. Among the bacterial genomes, this matches to *Tenacibaculum mesophilum* with BBS=776. This has no match in the ’nt’ database when the search is constrained to bacteria, demonstrating that all bacterial genomes are not included in the ’nt’ database.

### Fungal transcripts

Fungal transcripts (BBS=150, annoEnsFungi.anno.sort in Dataset1, n=795) reveal several commonly occuring genera, detected in small traces in different tissues (Table 6). The exact species can not be identified using the available data, since these genera incorporate many species. http://www.speciesfungorum.org/Names/names.asp provides the enumeration of fungal species. For example, *Mycosphaerella*, a genus of ascomycotaes, is one of the largest genus of plant pathogen fungi with more than 1500 species [32]. Similarly, *Pyrenophora* has ~160 species, several of them pathogenic [33]

**Table 6:** Fungal transcripts: These are sorted based on BLAST bitscore (BBS). Although, the best match specifies a genus and the species, this identification needs validation. The counts in the different tissues are lesser than those of viral and bacterial transcripts. STI: STIGMA, STA: STAMEN.

| TRID | Accid | Description | BBS | CORM | TEPAL | LEAF | STI | STA |
| --- | --- | --- | --- | --- | --- | --- | --- | --- |
| CSTC021516 | XM_002619271.1 | *Clavispora lusitaniae* | 1897 | 2 | 581 | 0 | 31 | 11 |
| CSTC086633 | XM_013574385.1 | *Aureobasidium namibiae* | 1306 | 4 | 345 | 5 | 398 | 25 |
| CSTC072893 | XM_001551736.1 | *Botryotinia fuckeliana* | 1290 | 5 | 360 | 2 | 57 | 1 |
| CSTC101026 | DQ677898.1 | *Cladosporium cladosporioides* | 1210 | 1 | 142 | 1 | 64 | 8 |
| CSTC023760 | XM_001930596.1 | *Pyrenophora tritici-repentis* | 1051 | 4 | 221 | 0 | 92 | 2 |
| CSTC064623 | XM_003856981.1 | *Mycosphaerella graminicola* | 1083 | 0 | 203 | 1 | 105 | 4 |
| CSTC026015 | X81860.1 | *Cladosporium herbarum* | 1066 | 4 | 175 | 1 | 103 | 8 |
| CSTC016711 | XM_016737515.1 | *Penicillium expansum* | 885 | 3 | 99 | 0 | 48 | 0 |
| CSTC041979 | XM_003854273.1 | *Mycosphaerella graminicola* | 824 | 2 | 126 | 0 | 101 | 0 |

### Caveats and advantages of ORF based annotation

Annotation of transcripts can also be done by analyzing ORFs. Although the nucleotide sequence of CSTC042320 has no match in the complete ’nt’ database, it encodes a 479 long ORF homologous (57% identity) to a cytochrome P450 protein from pineapple (*Ananas comosus*). The saffron transcriptome has been annotated accordingly (Saffron functional annotation [7]), and has significant match in the online interface to the saffron genome (http://www.crocusgenome.org/?pageid=20). However, such functional annotation of proteins in the absence of a genome is subject to errors due to the existence of significantly homologous proteins in organisms from different branches of life, like fungi, plants and viruses. CSTC018609 is one such mis-annotated transcripts, that encodes an 213 long ORF, homologous to a heat shock protein from the fungi *Cladosporium cladosporioides* with 99% identity, but is annotated as a plant gene (Saffron functional annotation [7]). The nucleotide sequence of CSTC018609 also matches to the genome of the fungi *Cladosporium cladosporioides* (Accid:HQ693482.1) with BBS=1094, 97% identity. Although the nucleotide sequence also matches to a plant (*Nicotiana tabacum*, Accid:AY372070.1) with lesser significance (87% identity), it has no significant match through the online interface provided. Interestingly, the ORF of this gene is also homologous (76% identity) to a viral HSP70 protein (*Micromonas pusilla* virus, Accid:AET43623.1).

ORF based annotation provides coverage in some cases where nucleotide homology is not detected. The viral protein RefSeq database identified additional ∼20 putative viral transcripts (see viral.anno.protein.txt in Dataset1) not detected through the nucleotide comparison. One reason for such an exclusion is the smaller size of the local viral nucleotide database sequence, which does not include all strains. CSTC027157 has significant matches among the full ’nt’ virus database (bean common mosaic virus complete genome, isolate PV 0915, Accid:HG792064.1). However, some sequences (CSTC004726) have no match even in the complete ’nt’ database, although they encode ORFs with significant matches to the local viral protein database. CSTC004726 encodes a polyprotein (Accid:NP 660175.1, Evalue=9e-37) from bean common mosaic necrosis virus (Evalue=9e-37). Thus, an annotation methodology should include comparisons to both nucleotide and amino acid sequences.

## Conclusions and future work

Previous application of the YeATS suite to genomes [9, 10] and metagenomes [11] required a BLAST to the complete ’nt’ and ’nr’ databases. In the current work, smaller, yet comprehensive databases have been created to accelerate the identification of the saffron metagenome from the transcriptome obtained from saffron in Jammu and Kashmir [7]. Transcripts from the soybean mosaic virus was found abundantly expressed in all five tissues analyzed. Although, these viruses have been isolated from these plants (and the related species *Iris*) previously [18–22], recent studies have shown the threat posed by these viruses to saffron cultivation is seriously underestimated. Several bacterial and fungal genera incorporating known pathogen species have also been identified.

A search through the complete set of sequences known (through a program like BLAST) is ideal, but not time efficient. The search can be accelerated by creating smaller, but comprehensive, sequence databases. Such a ’divide and conquer’ strategy needs to account for the existence of homologous nucleotide sequences (and proteins) across the different trees of life. There are two methods for annotating a transcript – (a) through the nucleotide sequence or (b) through the amino acid sequence (assuming it is not a non-coding transcript). Annotation based on ORFs identifies transcripts that have no nucleotide sequence homology in the entire ’nt’ database, which occurs due to the redundancy in the codon table. Finally, this work also highlights that the standard practice of excluding transcripts homologous to ribosomal RNA might exclude bona fide bacterial transcripts, in the absence of comparative analysis of the homology to different databases. While the current strategy of creating smaller, comprehensive subsets accelerates the annotation time as compared to YeATS, the existing flow still uses command-line BLAST to query the entire transcriptome with the sequence databases. A kmer based approach could accelerate this further, and will be added in a future release [34].

## Acknowledgements

I gratefully acknowledge Mridul Bhattacharjee and Nitin Salaye for logistic support.

## References

1 Escribano J, Alonso GL, Coca-Prados M, Fernández JA (1996) Crocin, safranal and picrocrocin from saffron (crocus sativus l.) inhibit the growth of human cancer cells in vitro. Cancer letters 100: 23–30.

2 Abdullaev FI (2002) Cancer chemopreventive and tumoricidal properties of saffron (crocus sativus l.). Experimental biology and medicine 227: 20–25.

3 Lopresti AL, Drummond PD (2014) Saffron (crocus sativus) for depression: a systematic review of clinical studies and examination of underlying antidepressant mechanisms of action. Human Psychopharmacology: Clinical and Experimental 29: 517–527.

4 Hausenblas HA, Saha D, Dubyak PJ, Anton SD (2013) Saffron (crocus sativus l.) and major depressive disorder: a meta-analysis of randomized clinical trials. Journal of integrative medicine 11: 377–383.

5 Wang Z, Gerstein M, Snyder M (2009) RNA-seq: a revolutionary tool for transcriptomics. Nature Reviews Genetics 10: 57–63.

6 Flintoft L (2008) Transcriptomics: digging deep with RNA-seq. Nature Reviews Genetics 9: 568–568.

7 Jain M, Srivastava PL, Verma M, Ghangal R, Garg R (2016) De novo transcriptome assembly and comprehensive expression profiling in crocus sativus to gain insights into apocarotenoid biosynthesis. Scientific reports 6.

8 Baba SA, Mohiuddin T, Basu S, Swarnkar MK, Malik AH, et al. (2015) Comprehensive transcriptome analysis of crocus sativus for discovery and expression of genes involved in apocarotenoid biosynthesis. BMC genomics 16: 1.

9 Chakraborty S, Britton M, Wegrzyn J, Butterfield T, Martinez-Garcia PJ, et al. (2015). YeATS-a tool suite for analyzing RNA-seq derived transcriptome identifies a highly transcribed putative extensin in heartwood/sapwood transition zone in black walnut.

10 Martínez-García PJ, Crepeau MW, Puiu D, Gonzalez-Ibeas D, Whalen J, et al. (2016) The walnut (juglans regia) genome sequence reveals diversity in genes coding for the biosynthesis of nonstructural polyphenols. The Plant Journal.

11 Chakraborty S, Britton M, Mart´ınez-Garc´ıa P, Dandekar AM (2016) Deep RNA-seq profile reveals biodiversity, plant–microbe interactions and a large family of NBS-LRR resistance genes in walnut (juglans regia) tissues. AMB Express 6: 1.

12 Simpson JT, Wong K, Jackman SD, Schein JE, Jones SJ, et al. (2009) Abyss: a parallel assembler for short read sequence data. Genome research 19: 1117–1123.

13 Koonin EV, Dolja VV, Morris TJ (1993) Evolution and taxonomy of positive-strand rna viruses: implications of comparative analysis of amino acid sequences. Critical reviews in biochemistry and molecular biology 28: 375–430.

14 Byamukama E, Wegulo SN, Tatineni S, Hein G, Graybosch R, et al. (2014) Quantification of yield loss caused by triticum mosaic virus and wheat streak mosaic virus in winter wheat under field conditions. Plant Disease 98: 127–133.

15 Margulis L (1996) Archaeal-eubacterial mergers in the origin of eukarya: phylogenetic classification of life. Proceedings of the national academy of sciences 93: 1071–1076.

16 Yates A, Akanni W, Amode MR, Barrell D, Billis K, et al. (2016) Ensembl 2016. Nucleic acids research 44: D710–D716.

17 Camacho C, Madden T, Ma N, Tao T, Agarwala R, et al. (2013) BLAST Command Line Applications User Manual.

18 Russo M, Martelli G, Cresti M, Ciampolini F (1979) Bean yellow mosaic virus in saffron/il virus del mosaico giallo del fagiolo in zafferano. Phytopathologia Mediterranea : 189–191.

19 Miglino R, Jodlowska A, Van Schadewijk A (2005) First report of narcissus mosaic virus infecting crocus spp. cultivars in the netherlands. Plant Disease 89: 342–342.

20 Yan S, Qin Z, Jin L, Chen J (2010) A new isolate of iris severe mosaic virus causing yellow mosaic in iris ensata thunb. Journal of nanoscience and nanotechnology 10: 726–730.

21 Nateqi M, Habibi MK, Dizadji A, Parizad S (2015) Detection and molecular characterization of the iris severe mosaic virus-ir isolate from iran. Journal of Plant Protection Research 55: 235–240.

22 Wylie SJ, Li H, Liu J, Jones MG (2014) First report of narcissus mosaic virus from australia and from iris. Australasian Plant Disease Notes 9: 1–2.

23 Caiola MG, Faoro F (2011) Latent potyvirus infections in crocus sativus artwrightianus: an under-estimated problem in saffron? Phytopathologia Mediterranea 50: 175–182.

24 Doolittle WF (1998) A paradigm gets shifty. Nature 392: 15–16.

25 Švec P, Vancanneyt M, Sedláček I, Naser SM, Snauwaert C, et al. (2006) Enterococcus silesiacus sp. nov. and enterococcus termitis sp. nov. International journal of systematic and evolutionary microbiology 56: 577–581.

26 Czekaj T, Ciszewski M, Szewczyk EM (2015) Staphylococcus haemolyticus–an emerging threat in the twilight of the antibiotics age. Microbiology 161: 2061–2068.

27 Grisham M, Pan YB, Richard Jr E (2007) Early detection of leifsonia xyli subsp. xyli in sugarcane leaves by real-time polymerase chain reaction. Plant Disease 91: 430–434.

28 Young A, Petrasovits L, Croft B, Gillings M, Brumbley S (2006) Genetic uniformity of international isolates of leifsonia xyli subsp. xyli, causal agent of ratoon stunting disease of sugarcane. Australasian Plant Pathology 35: 503–511.

29 Jean S, Lee WS, Chen F, Ou T, Hsueh P (2014) Elizabethkingia meningoseptica: an important emerging pathogen causing healthcare-associated infections. Journal of Hospital Infection 86: 244–249.

30 Balm M, Salmon S, Jureen R, Teo C, Mahdi R, et al. (2013) Bad design, bad practices, bad bugs: frustrations in controlling an outbreak of elizabethkingia meningoseptica in intensive care units. Journal of Hospital Infection 85: 134–140.

31 Shinha T, Ahuja R (2015) Bacteremia due to elizabethkingia meningoseptica. IDCases 2: 13–15.

32 Crous P, Braun U, Groenewald J (2007) Mycosphaerella is polyphyletic. Studies in Mycology 58: 1–32.

33 Day J, Gietz RD, Rampitsch C (2015) Proteome changes induced by pyrenophora tritici-repentis toxa in both insensitive and sensitive wheat indicate senescence-like signaling. Proteome science 13: 1.

34 Compeau PE, Pevzner PA, Tesler G (2011) How to apply de bruijn graphs to genome assembly. Nature biotechnology 29: 987–991.

